## Supplemental Figure S1-S4 for "ScRNA-seq of Diverse Pheochromocytoma Patients Reveals Distinct Microenvironment Characteristics and Supports an Informative Molecular Classification System"

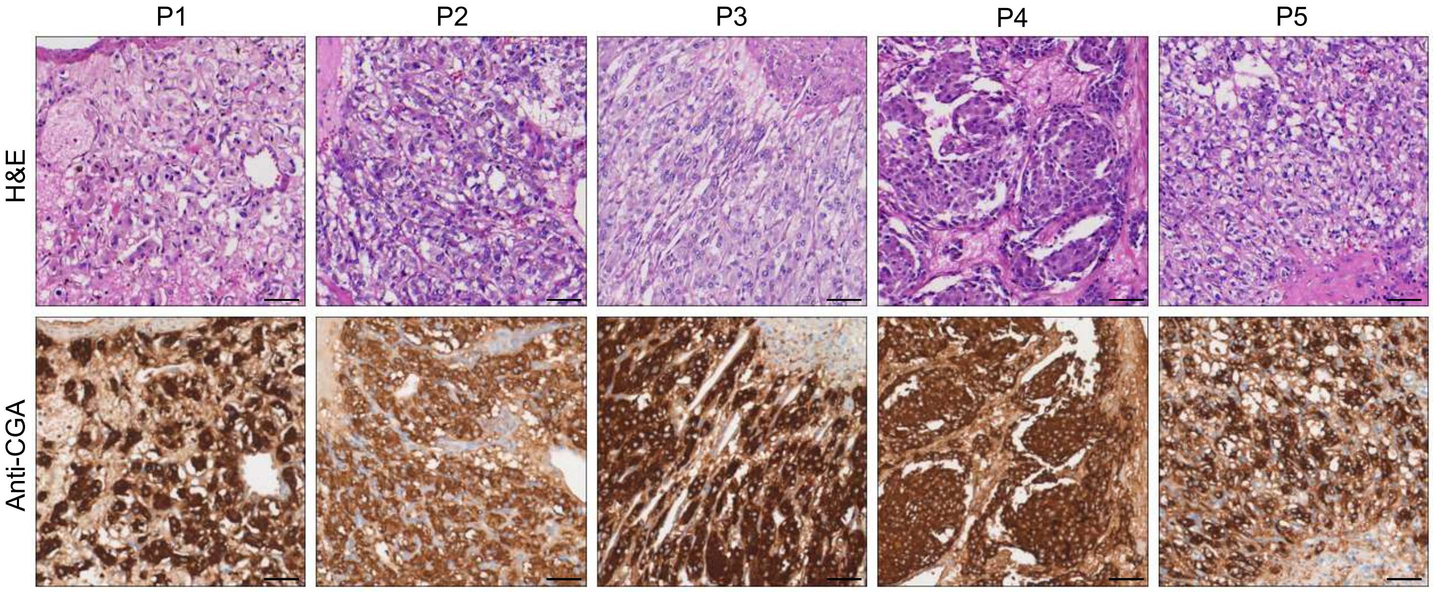


**Supplemental Figure S1. Hematoxylin-eosin Staining and Immunohistochemistry Staining of CGA Marker in Formalin-fixed Paraffin-embedded PCC Tissue Sections Matched to scRNA-seq Specimens.** Scale bar, 100 μm.

#### Supplemental Figure S2

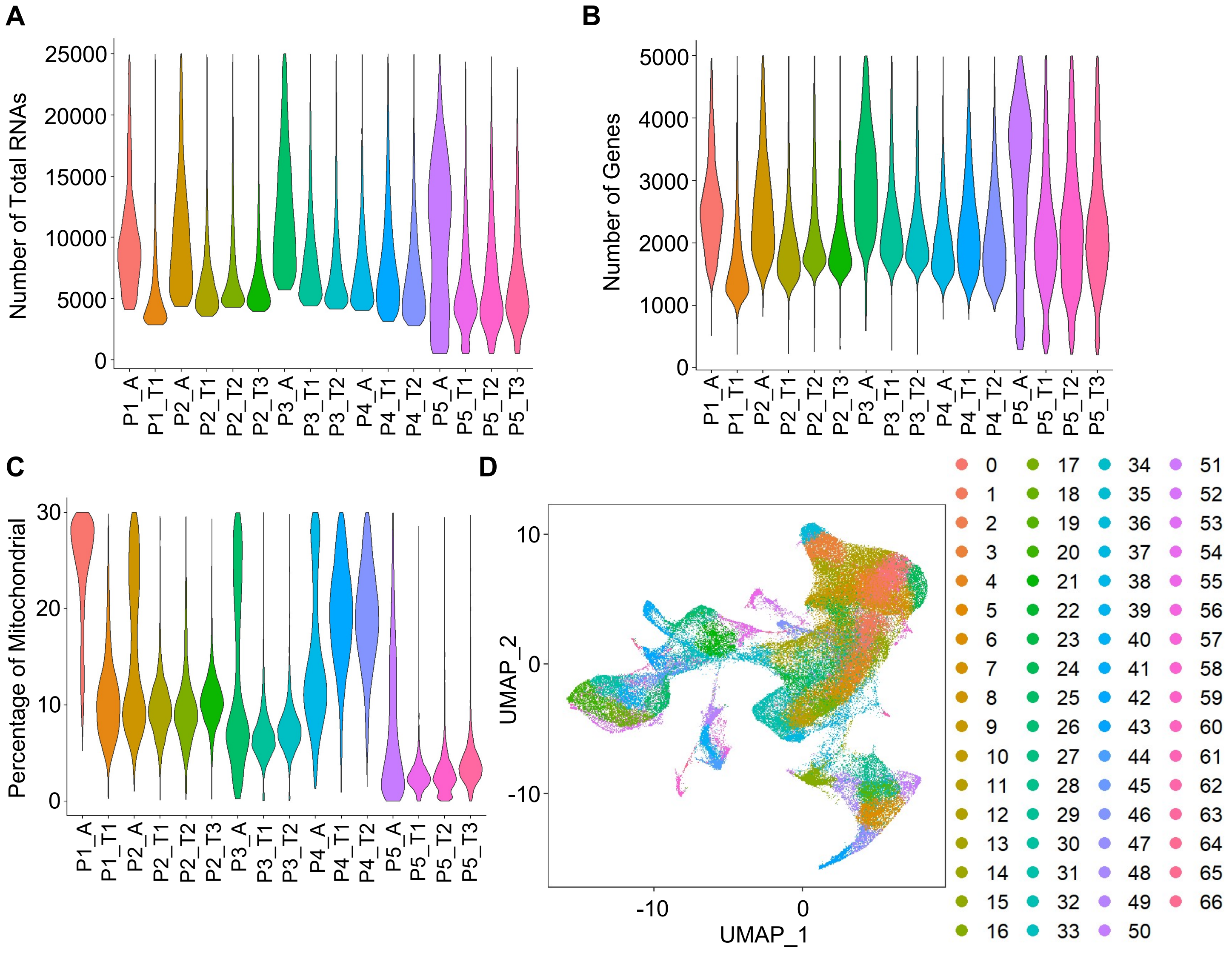

### Supplemental Figure S2

**E**

**Correlation among Clusters**

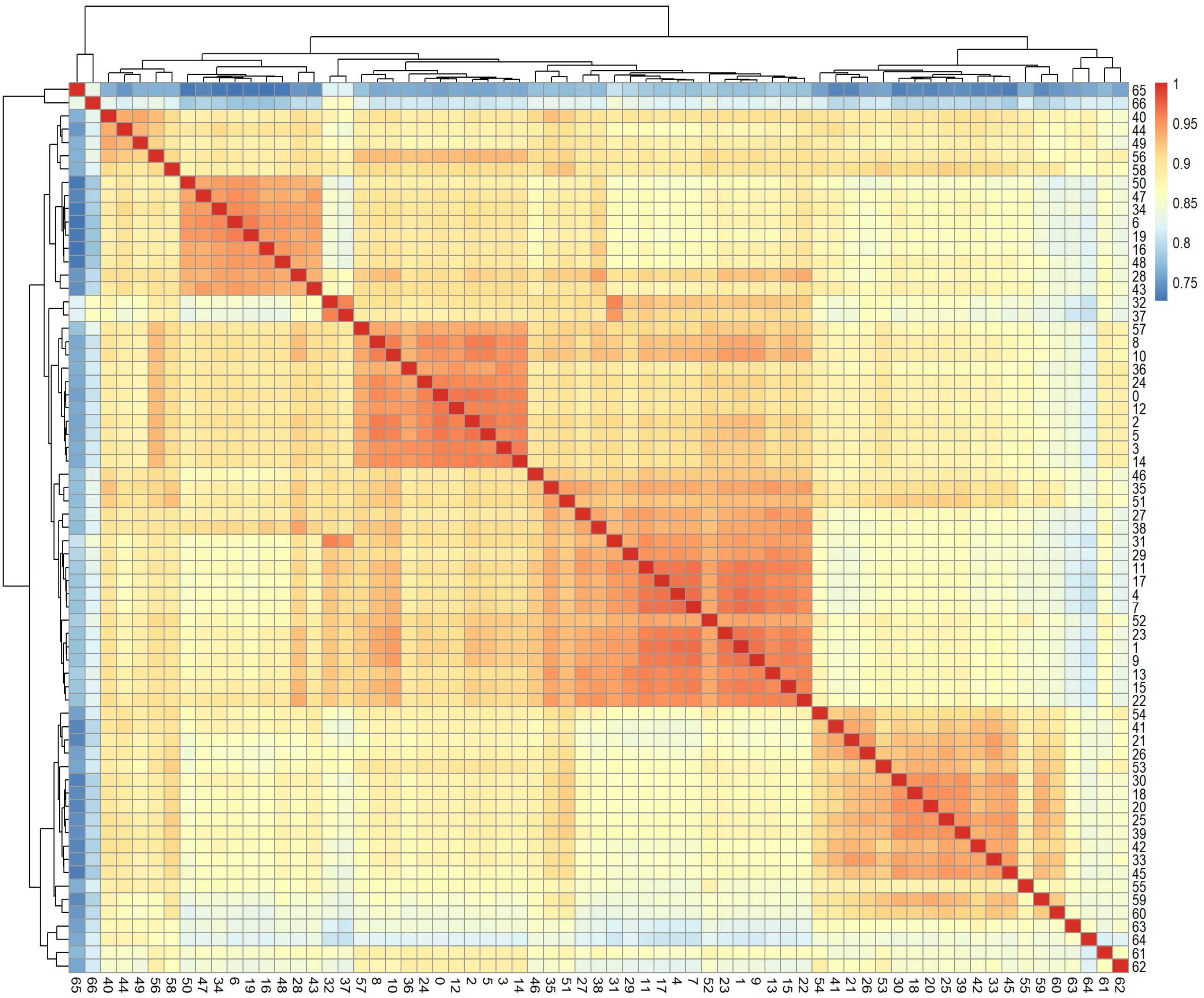


**Supplemental Figure S2. Quality Control and Cell Clustering of scRNA-Seq Data.** (A, B, C) Violin plots showing the number of total RNAs (A), the number of genes (B), and the percentage of mitochondrial (mito) genes (C) for cells from 16 specimens. (D) UMAP plots of cells colored by cell clusters. (E) Heatmap plotting the correlation coefficient between cell clusters. The color keys from blue to red indicate the correlation coefficient from low to high.

#### Supplemental Figure S3

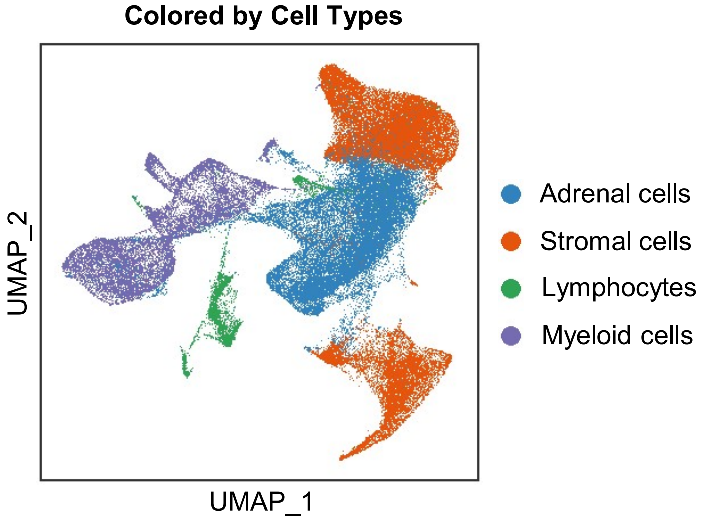


**Supplemental Figure S3. Integration Analysis across 5 PCC Patients Revealing the Cell Type Composition of the PCC Microenvironment.** UMAP plot depicting the distribution of adrenal cells, stromal cells, and immune cells (including lymphocytes and myeloid cells) within the PCC microenvironment.

#### Supplemental Figure S4

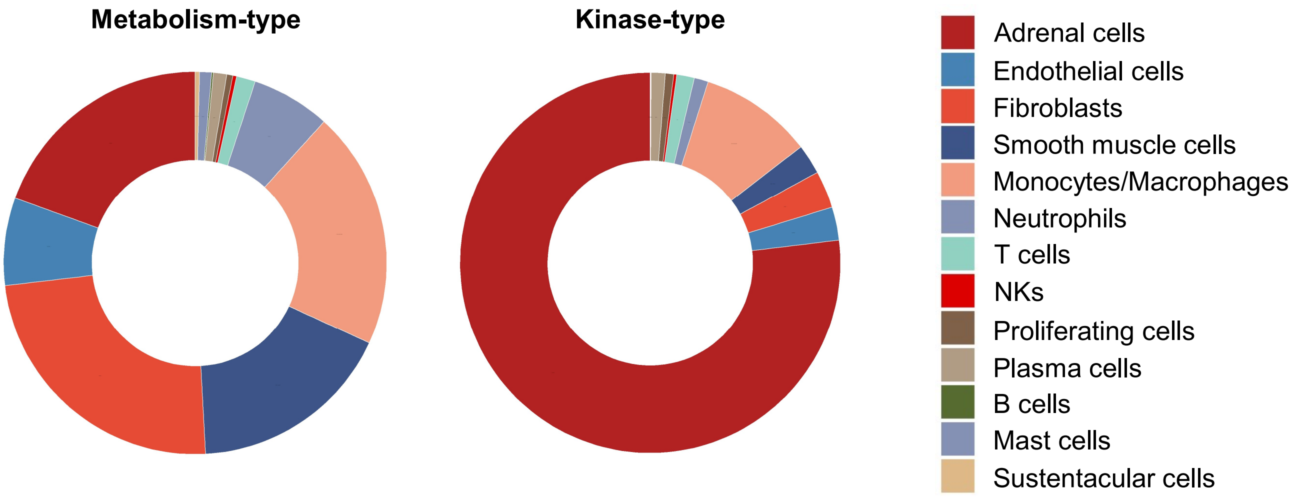


**Supplemental Figure S4. The Frequency Distribution of Cell Types within the Microenvironment of Metabolism-type and Kinase-type PCC Patients.**
