## Supplemental Table S1-S3 for "ScRNA-seq of Diverse Pheochromocytoma Patients Reveals Distinct Microenvironment Characteristics and Supports an Informative Molecular Classification System"

**Supplemental Table S1: Somatic and Germline Mutations in 5 PCC Patients Detected by WES**

| | P1 | P2 | P3 | P4 | P5 |
|---|---|---|---|---|---|
| **Somatic Mutations (Mutation Rates)** | JAK2 (3.2%) | ARHGEF39 (5.33%) | KMT2D (6.21%) | MST1 (2.68%) | REV3L (2.74%) |
| | NQO1 (2.97%) | METTL4 (5.09%) | CHEK2 (1.46%) | RPA1 (1.69%) | RYR2 (2.3%) |
| | CDH18 (2.83%) | PAK1IP1 (5.07%) | KDM6A (1.05%) | APCDD1 (1.52%) | KMT5A (2.19%) |
| | KCNT2 (2.78%) | IGSF3 (3.2%) | CSDE1 (1.05%) | SMARCA4 (1.14%) | |
| | HCN1 (2.48%) | MST1 (0.85%) | | STK11 (1.05%) | |
| | INSRR (2.02%) | | | | |
| **Germline Mutation (Mutation Site)** | N/A | N/A | N/A | N/A | VHL (c.499C>T) |

**Supplemental Table S2: PASS Scores of Collected Tumor Tissues**

| | P1_T1 | P2_T1 | P2_T2 | P2_T3 | P3_T1 | P3_T2 | P4_T1 | P4_T2 | P5_T1 | P5_T2 | P5_T3 |
|---|---|---|---|---|---|---|---|---|---|---|---|
| **Large nest/diffuse growth >10% of tumor volume** | 2 | 2 | 2 | 2 | 2 | 2 | 2 | 2 | 2 | 2 | 2 |
| **Central or confluent tumor necrosis** | 0 | 0 | 0 | 0 | 0 | 0 | 0 | 0 | 0 | 0 | 0 |
| **High cellularity** | 2 | 0 | 0 | 2 | 0 | 2 | 2 | 2 | 0 | 2 | 0 |
| **Cellular monotony** | 0 | 0 | 2 | 2 | 0 | 2 | 0 | 0 | 0 | 2 | 0 |
| **Tumor cell spindling** | 0 | 0 | 0 | 0 | 0 | 0 | 0 | 0 | 0 | 0 | 0 |
| **Mitotic figures >3/10 high power field** | 0 | 0 | 0 | 0 | 0 | 2 | 2 | 0 | 0 | 0 | 2 |
| **Atypical mitotic figures** | 0 | 0 | 0 | 0 | 0 | 0 | 0 | 0 | 0 | 0 | 0 |
| **Extension into adipose tissue** | 0 | 0 | 0 | 0 | 0 | 0 | 0 | 0 | 0 | 0 | 0 |
| **Vascular invasion** | 0 | 0 | 0 | 1 | 0 | 0 | 1 | 0 | 0 | 1 | 1 |
| **Capsular invasion** | 0 | 0 | 0 | 0 | 0 | 0 | 1 | 0 | 0 | 1 | 0 |
| **Profound nuclear pleomorphism** | 0 | 0 | 0 | 0 | 0 | 0 | 0 | 0 | 0 | 0 | 0 |
| **Nuclear hyperchromasia** | 1 | 0 | 0 | 0 | 1 | 1 | 1 | 0 | 0 | 1 | 0 |
| **Total Score** | 5 | 2 | 4 | 7 | 3 | 9 | 9 | 4 | 2 | 9 | 5 |

**Supplemental Table S3: Clinical Information of 5 PCC Patients**

| | Metabolism-type PCC | | | | Kinase-type PCC |
|---|---|---|---|---|---|
| | **P1** | **P2** | **P3** | **P5** | **P4** |
| **Tumor Size ($cm^3$)** | 5 × 3.8 × 3.7 | 3 × 3 × 2.5 | 5.5 × 5.5 × 4 | 3.2 × 3 × 1.7 | 6 × 5 × 4 |
| **Blood Pressure (mmHg)** | 160/110 | 120/77 | 148/108 | 170/110 | 200/120 |
| **Symptom** | Hypertension<br>Headache<br>Palpitation | Asymptomatic | Hypertension<br>Dizziness<br>Weakness<br>Fever | Hypertension<br>Dizziness<br>Weakness | Hypertension<br>Headache<br>Palpitation<br>Hyperhidrosis |
| **3-methoxytyramine (3-MT) (pmol/L)** | 46.2 | ＜24 | ＜24 | 37.9 | 53.7 |
| **Metanephrine (MN) (pmol/L)** | 145.7 | 120.2 | 1423.6 | 62.8 | 113.4 |
| **Normetanephrine (NMN) (pmol/L)** | 11928.5 | 5090.7 | 10486.6 | 2888.5 | 19215.8 |
